# The Dream and MEC NuRD Complexes reinforce SPR-5/MET-2 maternal reprogramming to maintain the germline-soma distinction

**DOI:** 10.1101/2025.07.23.666413

**Authors:** Sindy R. Chavez, Jazmin Dozier, Saahj P. Gosrani, Sandra K. Nguyen, Jovan S. Brockett, Sarah D. Blancher, Sydney L. Morgan-Benitez, Juan D. Rodriguez, Onur Birol, Monica N. Reeves, Karen L. Schmeichel, David J. Katz, Brandon S. Carpenter

## Abstract

The proper coordination of transcription factors, ATP dependent chromatin remodelers and histone modifications is essential for tissue specific gene expression, but how gene expression is regulated at these different levels is not well understood. In *C. elegans*, H3K4 methylation that is acquired in the germline is reprorgammed at fertilization by the H3K4me1/2 demethlyase SPR-5/LSD1/KDM1A and the H3K9 methyltransferase MET-2/SETDB1/KMT2E. SPR-5/MET-2 maternal reprogramming is required to help establish the germline-soma distinction and prevent developmental delay by preventing inherited H3K4 methylation from inappropriately maintaining germline gene expression in somatic tissues. To determine if the DREAM transcriptional repressor complex and the MEC NuRD ATP dependent nucleosome remodeling and histone deacetylase complex function to reinforce SPR-5/MET-2 maternal reprogamming, we asked if loss of these complexes affects the ectopic germline transcription and developmental delay in *spr-5; met-2* double mutants. We find that knocking down the DREAM or MEC NuRD complexes specifically exacerbates the developmental delay and ectopic expression of germline genes in the soma caused by loss of SPR-5 and MET-2. In addition, the DREAM and MEC NuRD complexes bind together at SPR-5/MET-2 reprogramming targets. These data suggest that the transcriptional repression of DREAM and the ATP dependent chromatin remodeling and deactylation activities of the MEC NuRD complex are required somatically to reinforce maternal histone reporgamming by SPR-5/MET-2. Thus, these data provide a novel example of how gene regulation is coordinated at multiple levels to maintain the germline-soma distinction and ensure proper development.

## INTRODUCTION

Gene transcription must be coordinately regulated to establish tissue specific gene expression and ensure normal development. To accomplish this, genes must be regulated at multiple levels, including by transcription factors, ATP dependent chromatin remodelers and histone modifications. However, how tissue specific gene expression is coordinately regulated at these different levels is not well understood.

In *C. elegans,* the specification of germline versus soma is partially established by the coordinate regulation of inherited histone modifications. During gametogenesis, sperm and oocyte-specific genes acquire transcription-coupled histone methylation which helps sustain germline gene expression by maintaining DNA access to transcription factors and RNA polymerase (Burton and Torres-Padilla, 2014; Hirabayashi and Gotoh, 2010; Jambhekar et al., 2019). Transcription acquired histone methylation can be inherited through mitotic cell divisions and between generations via both the sperm and oocyte. For this reason, the newly formed zygote must reset, or “reprogram”, the inherited histone methylation to prevent the inappropriate maintenance of germline gene expression (Gaydos et al., 2014; Jambhekar et al., 2019; Kaneshiro et al., 2019; Öst et al., 2014; Siklenka et al., 2015; Tabuchi et al., 2018; Zheng et al., 2016). To do this, maternally deposited histone modifying enzymes add and remove methyl groups on the N-terminal tails of histone core proteins to limit DNA accessibility and reset the chromatin landscape. For example, in *C. elegans* and mice, the histone demethylase SPR-5/LSD1, removes mono- and di-methylation of lysine 4 on histone H3 (H3K4me1/2) at fertilization to prevent the inappropriate maintenance of germline gene expression in the embryo (Ancelin et al., 2016; Kim et al., 2015a; Stewart et al., 2015; Wasson et al., 2016). SPR-5/LSD1 functions together with the H3K9me2 methyltransferase, MET-2/SETDB1, during reprogramming at fertilization and this reprogramming is also conserved in mice (Greer et al., 2014; Katz et al., 2009; Kerr et al., 2014; Kim et al., 2016). In *C. elegans,* progeny of mutants lacking both SPR-5 and MET-2 are developmentally delayed and exhibit maternal effect sterility (Carpenter et al., 2021; Greer et al., 2014; Kerr et al., 2014). SPR-5/MET-2 maternal reprogramming is antagonized by the H3K36 methyltransferase MES-4, which transgenerationally maintains H3K36 methylation that is acquired in the germline of the previous generation and inherited in the embryo of the progeny (Furuhashi et al., 2010; Rechtsteiner et al., 2010). Hereafter, the set of germline genes that are bound by MES-4 and in which H3K36 methylation is transgenerationally maintained will be referred to as MES-4 germline genes. In progeny lacking SPR-5/MET-2 maternal reprogramming, the MES-4 germline genes are inappropriately expressed in somatic tissues. This inappropriate somatic expression causes a severe developmental delay (Carpenter et al., 2021).

Because the specification of germline versus soma is partially established by the coordinate regulation of inherited histone modifications, it provides the perfect opportunity to investigate how the regulation of transcription factors and chromatin remodeling complexes coordinate with histone modifications to regulate tissue specific transcription. In *C. elegans,* loss of either the DREAM or MEC **Nu**cloesome **R**emo**d**elling (NuRD) complexes also leads to somatic expression of germline genes in the soma (Costello and Petrella, 2019; Cui et al., 2006; Gal et al., 2021; Goetsch et al., 2017; Petrella et al., 2011; Rechtsteiner et al., 2019; Unhavaithaya et al., 2002; Wang et al., 2005; Wu et al., 2012). As a result, we considered the possibility that these complexes may be functioning coordinately with SPR-5/MET-2 maternal reprogramming to help specify germline versus soma. The DREAM complex, which includes LIN-35 (homolog of human retinoblastoma (rb)protein), LIN-9, LIN-37, and LIN-52, is a transcriptional repressor complex most known for its roles in repressing cell-cycle reentry genes (Dick and Rubin, 2013; Lu and Horvitz, 1998). At high temperatures, loss of DREAM complex members lead to an early larval arrest at the L1 stage and this L1 arrest can be rescued by knocking down *mes-4* (Rechtsteiner et al., 2019).

The NuRD complex is thought to primarily function as a repressor. It accomplishes this via two different activities: 1) ATP-dependent nucleosome remodeling, which represses chromatin by limiting the spacing between nucleosomes, and 2) histone deactylation, which primarily tightens the association between nucleosomes and DNA (Xue et al., 1998). In *C. elegans,* the histone deacytlation NuRD-like MEC complex contains MEP-1, LET-418 and HDA-1(Passannante et al., 2010). Similar to loss of DREAM, loss of the MEC NuRD complex component MEP-1 in *C. elegans* causes germline gene expression in somatic tissues (Unhavaithaya et al., 2002; Wang et al., 2005). This leads to a developmental delay that phenocopies *spr-5; met-2* mutants and can be rescued by knocking down *mes-4* (Unhavaithaya et al., 2002).

Here, we examined the synergism between SPR-5/MET-2 reprogramming and the highly conserved DREAM and MEC NuRD complexes and demonstrate that loss of either the DREAM or MEC NuRD complex in *spr-5; met-2* mutants causes an L1 larval arrest that is more severe than the L2 larval delay that we observe in *spr-5; met-2* mutants. We also find that similar genes that are mis-regulated in the soma of *spr-5; met-2* are further exacerbated upon loss of either Dream or MEC NuRD complex members. Finally, using published ChIP-seq data sets we confirm that MEP-1 and LIN-35 directly bind genes the genes that are misregualted in the soma of *spr-5; met-2* mutants. Together, our findings suggest that SPR-5, MET-2, the Dream Complex and the MEC NuRD Complex cooperate to maintain the germline-soma distinction in *C. elegans*.

## MATERIALS AND METHODS

### Strains

All *Caenorhabditis elegans* strains were grown and maintained at 20°C under standard conditions, as previously described (Brenner, 1974). Strains used were: N2: Bristol wild type strain provided by the *Caenorhabditis* Genetics Center; the *C. elegans spr-5 (by101)(I)* strain was provided by R. Baumeister (Albert Ludwig University of Freiburg, Germany); MT13293: *met-2 (n4256)(III) strain* was provided by R. Horvitz (Massachusetts Institute of Technology, MA, USA); the *spr-5 (by101)(I)*/*tmC27[unc-75(tmls1239)](I); met-2 (n4256) (III*)/qC1 [qls26 (lag2::gfp+ rol-6(su1006))](III) strain was created to maintain *spr-5 (by101)(I); met-2 (n4256)(III)* double-mutant animals as balanced heterozygotes (Carpenter et al., 2021).

### Scoring developmental delay

*C. elegans* adult hermaphrodites were allowed to lay embryos for 2-4 hours and then removed in order to synchronize the development of progeny. Progeny were then imaged and scored for developmental progression at 72 hours after the synchronized lay.

### RNA sequencing and analysis

Total RNA was isolated using TRIzol reagent (Invitrogen) from 500-1000 starved L1 larvae born at 20°C overnight on unseeded NGM plates. Total RNA was sent to Georgia Genomics and Bioinformatics Core (University of Georgia, Athens, Georgia) for standard Poly-A RNA-seq services (Illumina Nextseq, 50bp paired-end reads). Raw sequencing reads were checked for quality using FastQC (Wingett and Andrews, 2018), filtered using Trimmomatic (Bolger et al., 2014), and remapped to the *C. elegans* transcriptome (ce11, WBcel235) using HISAT2 (Kim et al., 2015b). Read count by gene was obtained by FeatureCounts (Liao et al., 2014). Differentially expressed transcripts (significance threshold, Wald test, p-adj < 0.05 with a log2 fold change >1 for up regulated gene and <-1 for down regulated) were determined using DESEQ2 (v.2.11.40.2) (Love et al., 2014). Transcripts per million (TPM) values were calculated from raw data obtained from FeatureCounts output. Subsequent downstream analysis was performed using R with normalized counts and p-values from DESEQ2 (v.2.11.40.2). Heatmaps were produced using the ComplexHeatmap R Package (Gu et al., 2016). Data was scaled and hierarchical clustering was performed using the complete linkage algorithm. In the linkage algorithm, distance was measured by calculating pairwise distance. Volcano plots were produced using the EnhancedVolcano package (v.1.20.0). Raw and processed RNAseq files have been deposited into Gene Expression Omnibus (www.ncbi.nlm.nih.gov/geo) under accession code GSE303062 (*lin-35* RNAi RNAseq) and GSE303085 (*mep-1* RNAi RNAseq).

### ChIP analysis

LIN-35 and MEP-1 ChIP data were retrieved from the publicly available ENCODE datasets (LIN-35: https://www.encodeproject.org/experiments/ENCSR472HTK/; MEP-1: https://www.encodeproject.org/experiments/ENCSR048MPV/). Single and paired files for each replicate were used for downstream analysis. Quality control using FastQC was performed before and after adapter trimming using Trimmomatic to verify appropriate trimming. Reads were aligned to the ce11 *c. elegans* genome using Bowtie2. Peak calling was performed using macs2. To visualize peaks, bigWig files were generated from the ChIP data (bin size 20, normalized using RPKM). Heatmap matrices were prepared by combining score files (bigWig) and region files (BED) obtained from RNAseq datasets. Heatmaps were generated using the corresponding matrix.

### RNAi methods

RNAi by feeding was carried out using clones from the Ahringer library (Kamath and Ahringer, 2003). Feeding experiments were performed on RNAi plates (NGM plates containing 100 ug/ml ampicillin, 0.4mM IPTG, and 12.5ug/ml tetracycline). F0 worms were placed on RNAi plates as L2 larvae and then moved to fresh RNAi plates 48hrs later where they were allowed to lay embryos for 2-4 hrs. F0 worms were then removed from plates and sacrificed or placed on unseeded RNAi plates overnight so that starved L1 progeny could be isolated for RNAseq experiments. F1 progeny were scored 72hrs after the synchronized lay for L1 larval arrest. For each RNAi experiment, *pos-1* RNAi was used as a positive control. Each RNAi experiment reported here *pos-1* RNAi resulted in >95% embryonic lethality, indicating that RNAi plates were optimal.

### Differential interference contrast microscopy

Worms were immobilized in 0.1% levamisole and placed on a 2% agarose pad for Differential Interference Contrast (DIC) imaging at 10x magnification.

## RESULTS

### Loss of the DREAM or MEC NuRD complexes enhances the developmental delay of *spr-5; met-2* mutants

Previously we found that progeny of *spr-5; met-2* double mutants have a severe L2 developmental delay, with a small number of progeny becoming sterile adults approximately one week after being laid (Carpenter et al., 2021). To determine if the DREAM and MEC NuRD complexes are functioning in the SPR-5/MET-2 maternal reprogramming pathway, we asked whether knocking down DREAM and MEC NuRD complex components by RNA interference (RNAi) in *spr-5; met-2* mutants affects the severe L2 developmental delay. After 72 hours (hrs), all wild type (N2) worms fed empty vector control (L4440) RNAi reached adult stages (Figure 1A, K). Likewise, 72 hrs after hatching, 82% of progeny from *spr-5; met-2* mutants fed control RNAi proceeded past the L1 stage (Fig. 1B, K). But as we previously reported, almost all of these worms were delayed as L2 larvae (Carpenter et al., 2021). In controls, after 72 hrs 100% of progeny from wild type hermaphrodites fed *lin-35, lin-9, lin-37* or *lin-52* RNAi proceeded past the L1 stage (Fig. 1C, E, G, I, K), with most of these progeny reaching the adult stage. Thus, knockdown of the DREAM complex alone is not sufficient to result in developmental delay under these conditions. In contast, after 72 hrs, 100% of progeny from *spr-5; met-2* mutants fed RNAi against DREAM complex components *lin-35, lin-9, lin-37* or *lin-52* remain at the L1 larval stage (Fig. 1D, F, H, J, K), with almost all of these progeny being permanently arrested and never proceeding past the L1 stage. These data suggests that DREAM functions with SPR-5 and MET-2 to help development proceed normally.

**Figure 1.**
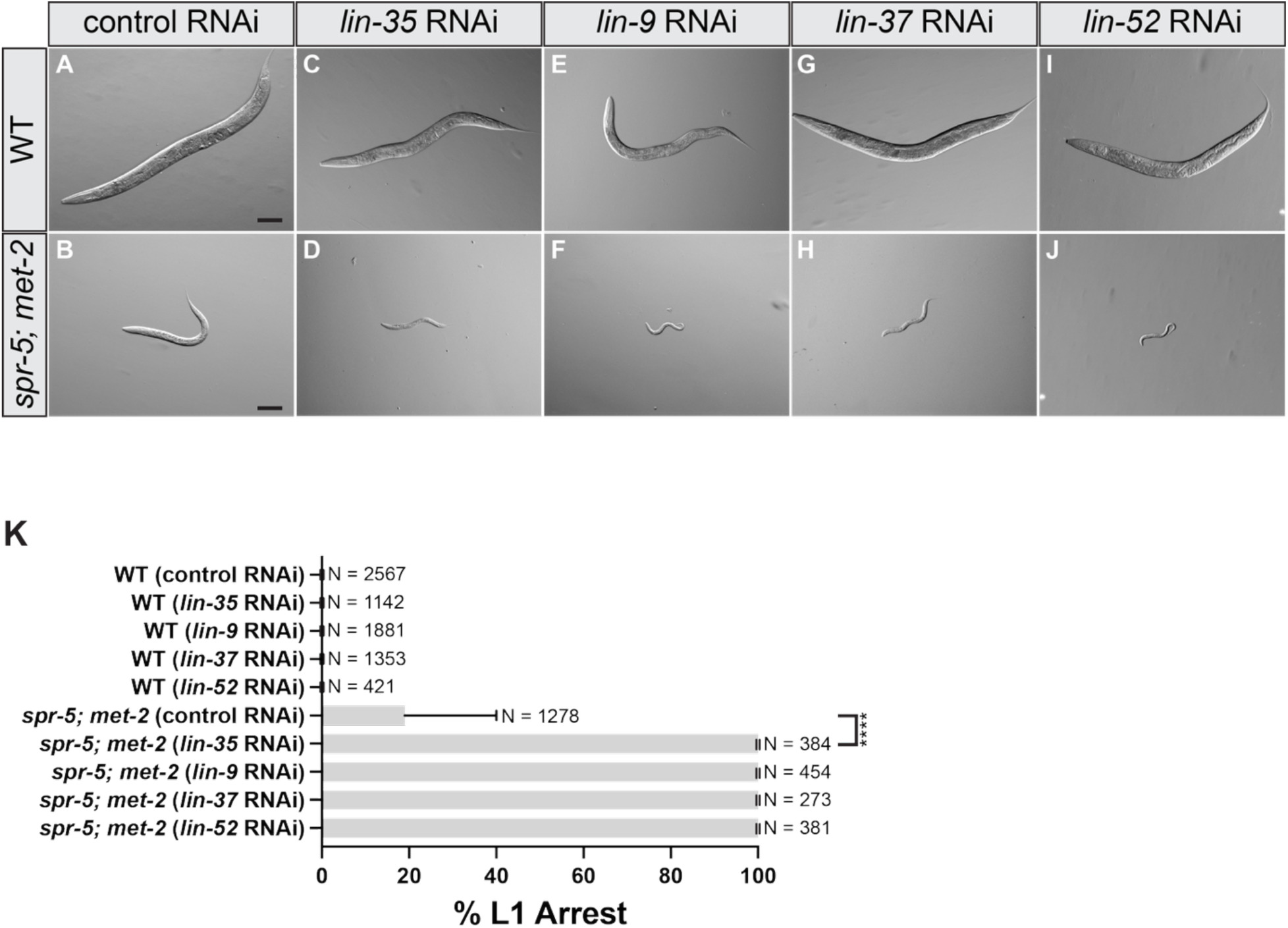
Knockdown of the DREAM complex causes L1 arrest in *spr-5; met-2* mutants. (A-J) 10x DIC (differential interference contrast) images taken of progeny 72 hours after synchronized lay. wild type (WT) fed either control (L4440) RNAi (A), *lin-35* RNAi (C), *lin-9* RNAi (E), *lin-37* RNAi (G), or *lin-52* RNAi (I) and *spr-5; met-2* mutants fed either control RNAi (B), *lin-35* RNAi (D), *lin-9* RNAi (F), *lin-37* RNAi (H), or *lin-52* RNAi (J). (K) Quantification of (A-D) scoring the percentage of L1 arrested progeny in wild type and *spr-5; met-2* mutants after 72 hours of synchronized lay. Data are mean±s.d. N=the total number of progeny from 3 or more replicates. (****) denotes P-value <0.0001 (two-tailed unpaired t-test). Scale bars: 100 μm.

To address the interaction between the MEC NuRD complex and SPR-5/MET-2 maternal reprogramming, we also performed RNAi against *mep-1* in *spr-5; met-2.* We chose to perform RNAi against *mep-1* because amongst the *C. elegans* NuRD-like complexes, MEP-1 is unique to the MEC NuRD complex (Passannante et al., 2010). Similar to what we observe with DREAM, knocking down the MEC NuRD complex complex components also exacerbated the developmental delay caused by the loss of SPR-5 and MET-2. After 72 hrs, all wild type progeny from hermaphprodites fed control RNAi reached adult stages (Figure 2A, E). Likewise, 98.8% of progeny from *spr-5; met-2* mutants fed control RNAi proceeded past the L1 stage (Figure 2B, E), but were subsequently delayed at the L2 stage. The 98.8% of *spr-5; met-2* mutants on control RNAi that proceeded past the L1 stage is higher than the 82% that we observed in the DREAM experiments (Figure 1K). It is possible that the remaining 18% of *spr-5; met-2* progeny in the DREAM experiments that arrested as L1s may be caused by a transgenerational build-up of H3K4me2 due to maintaining the balanced *spr-5; met-2* mutant strain for multiple generations prior to selecting double homozygous mutants. As a control, 99.9% of progeny from wild type hermaphprodites fed *mep-*1 RNAi proceeded past the L1 stage, with most of these progeny reaching the adult stage after 72 hours (Figure 2C, E). This finding suggests that knocking down the MEC NuRD complex alone is not sufficient to cause developmental delay under these conditions. In contrast, 91% of progeny from *spr-5; met-2* mutants fed *mep-1* RNAi remain at the L1 larval stage after 72 hrs (Figure 2D, E). As with DREAM, almost all of these progeny are permanently arrested and do not proceed past the L1 stage. Taken together, these results suggest that the DREAM complex and the MEC NuRD complex function together with SPR-5/MET-2 maternal reprogramming to ensure normal development.

**Figure 2.**
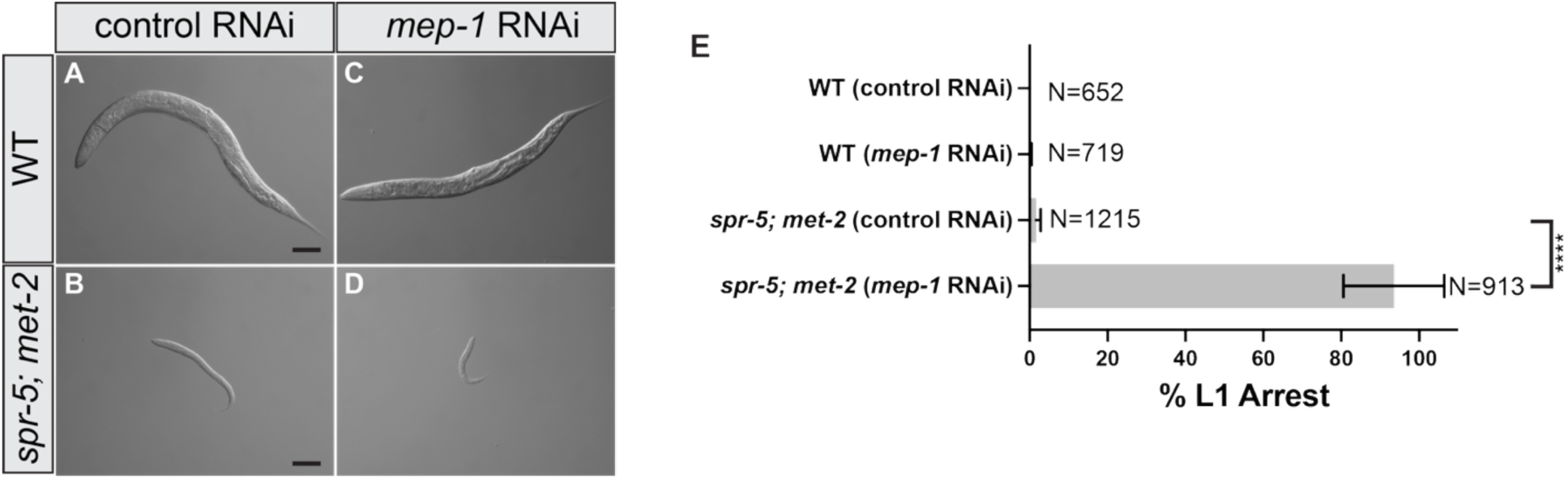
Knockdown of MEP-1 causes L1 arrest in *spr-5; met-2* mutants. (A-D) 10x DIC (differential interference contrast) images of progeny 72 hours after synchronized lay. wild type (WT) fed either control RNAi (A) or *mep-1* RNAi (C) and *spr-5; met-2* mutants fed either control RNAi (B) or *mep-1* RNAi (D). (E) Quantification of (A-D) scoring the percentage of L1 arrested progeny in wild type and *spr-5; met-2* mutants after 72 hours of synchronized lay. Data are mean±s.d.. N=the total number of progeny from 3 or more replicates. (****) denotes P-value <0.0001 (two-tailed unpaired t-test). Scale bars: 100 μm.

### Loss of the DREAM or MEC NuRD complexes specifically exacerbates *spr-5; met-2* gene expression changes

Since knocking down both DREAM and MEC NuRD complex members exacerbates the *spr-5; met-2* developmental delay, it raised the possibility that the DREAM and MEC NuRD complexes may specifically reinforce SPR-5/MET-2 maternal reprogramming. To determine if these functional interactions are specific, we performed RNA sequencing (RNAseq) on *spr-5; met-2* mutants fed RNAi against either DREAM or MEC NuRD complex components. Previously, we found that the developmental delay of *spr-5; met-2* mutants is caused by the expression of germline genes in somatic tissues (Carpenter et al., 2021). If the DREAM and MEC NuRD complexes are functioning to reinforce SPR-5/MET-2 maternal reprogramming, we would expect the ectopic expression of germline genes in the somatic tissues of *spr-5; met-2* mutants to be specifically exacerbated. To address this possibility, we performed RNAseq on starved L1 larvae. Starved L1 larvae have 550 somatic cells and only two germ cells, which have not begun proliferating or initiated germline expression. As a result, performing the RNAseq analysis on starved L1 larvae enables us to examine the ectopic expression of germline genes in somatic tissues.

To determine whether the ectopic expression of genes in *spr-5; met-2* mutants is exacerbated by the loss of the DREAM complex, we performed RNAseq on wild type L1 larvae compared to *spr-5; met-2* L1 larvae from hermaphrodites fed either control or *lin-35* RNAi (Supplementary Fig. 1). Compared to wild type, we identified 1,806 genes that are significantly up regulated in *spr-5; met-2* mutants with an average log2 fold change of 2.5 (Fig. 3A). Upon knockdown of *lin-35* in *spr-5; met-2* mutants, the average log2 fold change of these 1806 genes increases from 2.5 to 3.0 (Fig. 3A). This finding suggests that the DREAM complex reinforces SPR-5/MET-2 gene repression. As a control, we also compared the gene expression changes induced by RNAi of *lin-35* alone. Although there are some genes significantly up regulated by RNAi of *lin-35* alone, the average log2 fold change of the 1,806 genes up regulated in *spr-5; met-2* mutants compared to wild type in *lin-35* RNAi alone is only 0.3 (Fig. 3A). This suggests that the loss of the DREAM complex alone is generally not sufficient to upregulate genes that are ectopically expressed in the absence of SPR-5/MET-2 maternal reprogramming.

**Figure 3.**
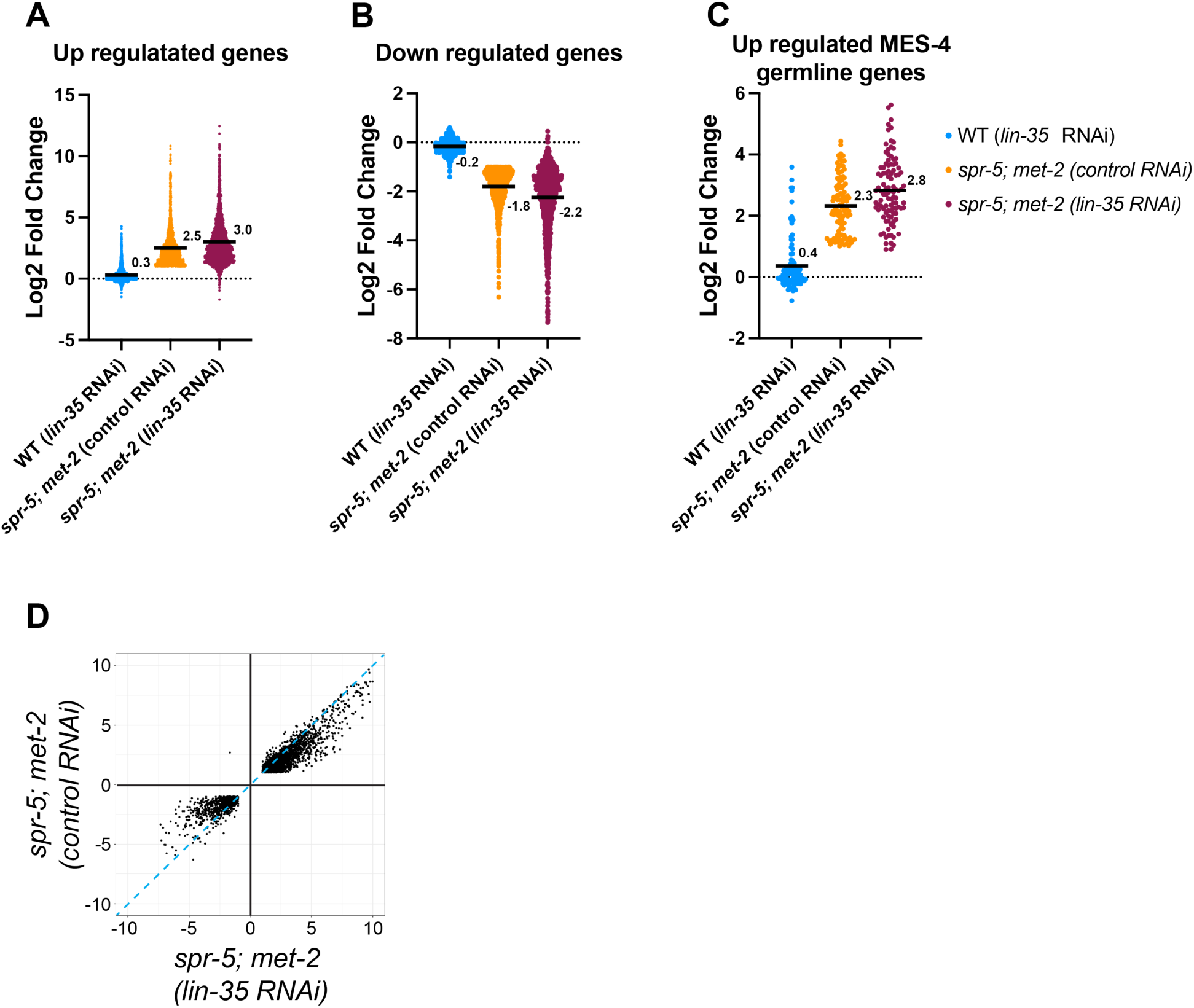
Knocking down *lin-35* further exascerbates differentially expressed genes in the soma of *spr-5; met-2* mutants. Scatter dot plots displaying the log2 fold change of the 1665 up-regulated genes (A), 752 down-regulated genes (B), and 101 up-regulated MES-4 germline genes (C) in *spr-5; met-2* mutants fed control RNAi (orange). The average log2 fold change of these same genes were examined in wild type (WT) hermaphrodites fed *lin-35* RNAi (blue) and *spr-5; met-2* mutants fed *lin-35* RNAi (maroon). (A-C) Numbers and solid black lines represent the mean log2 fold change. (D) Scatter plot displaying the correlation in log2 fold change of the all overlapping DEGs between spr-5; *met-2* mutants fed control RNAi and spr-5; *met-2* mutants *lin-35* RNAi L1 progeny. Dotted-blue line represents a one to one correlation in log2 fold change.

Knocking down *lin-35* also exacerbates genes that are down regulated in *spr-5; met-2* mutants. Of the 895 genes that are significantly down regulated in *spr-5; met-2* mutants compared to wild type, the average log2 fold change is -1.8 (Fig. 3B). When *lin-35* is knocked down in *spr-5; met-2* mutants, the average log2 fold change is -2.2 (Fig. 3B). This finding suggests that the DREAM complex is reinforcing SPR-5/MET-2 gene expression changes. Similar to what we observe with the up regulated genes, RNAi of *lin-35* alone causes a few of the 895 genes down regulated in *spr-5; met-2* mutants compared to wild type to be significantly down regulated. However, the average log2 fold change of these 895 down regulated genes in *lin-35* RNAi alone is only -0.2, demonstrating that loss of the DREAM complex alone is generally not sufficient to down regulate these genes (Fig. 3B).

To determine whether knocking down of *lin-35* specifically exacerbates the ectopic expression of MES-4 germline genes, we also examined the gene expression changes of the 176 germline genes targeted by MES-4 (Carpenter et al., 2021; Rechtsteiner et al., 2010).

There are 103 MES-4 germline genes that are significantly misregulated in *spr-5; met-2* mutants compared to wild type. Of these 103 MES-4 germline genes, 102 are up regulated with an average log2 fold change of 2.3, confirming that MES-4 germline genes are ectopically expressed in *spr-5; met-2* mutants (Fig. 3C). Upon knockdown of *lin-35* in *spr-5; met-2* mutants, this log2 fold change increases from 2.3 to 2.8 (Fig. 3C). These data suggest that the DREAM complex is reinforcing SPR-5/MET-2 repression of MES-4 germline genes in the soma. As a control, we also examined the expression of the MES-4 germline genes upon RNAi of *lin-35* alone. There are some MES-4 germline genes that are significantly up regulated by RNAi of *lin-35* alone. However, the average log2 fold change of the MES-4 germline genes in wild type animals fed *lin-35* RNAi is only 0.4, suggesting that loss of the DREAM complex alone is generally not sufficient to casue ectopic expression of MES-4 germline genes (Fig. 3C).

To examine the overall specificity of the interaction between DREAM and SPR-5/MET-2 reprogramming, we also examined the gene expression changes of all 2,389 genes that were changed in both *spr-5; met-2* mutants and *spr-5; met-2* mutants fed *lin-35* RNAi.

99.5% were changed in the same direction and of these 78.1% were exacerbated in *spr-5; met-2* mutants fed *lin-35* RNAi compared to *spr-5; met-2* mutants fed control RNAi. Only one gene was in an anti-correlated quadrant (Fig. 3D). Overall, these results suggest that the DREAM complex is functioning specifically to reinforce genes normally repressed by SPR-5 and MET-2.

To determine whether the ectopic expression of germline genes is exacerbated by the loss of the MEC NuRD complex, we also performed RNAseq on starved wild type L1 larvae compared to starved *spr-5; met-2* L1 larvae from hermaphrodites fed either control or *mep-1* RNAi (Supplementary Fig. 2). Compared to wild type, we indentified 1,665 genes that are significantly up regulated in *spr-5; met-2* mutants with an average log2 fold change of 2.5 (Fig. 4A). Knocking down *mep-1* in *spr-5; met-2* mutants caused the log2 fold change of these 1,665 genes to increase from 2.5 to 3.2, suggesting that the MEC NuRD complex is reinforcing SPR-5/MET-2 gene repression (Fig. 4A). Knocking down *mep-1* also exacerbates genes that are down regulated in *spr-5; met-2* mutants compared to wild type animals. Of the 752 genes that are significantly down regulated in *spr-5; met-2* mutants compared to wild type, the average log2 fold change is -1.6 (Fig. 4B). When *mep-1* is knocked down in *spr-5;met-2* mutants, the average log2 fold is -2.7, suggesting that the MEC NuRD complex is reinforcing SPR-5/MET-2 gene expression changes (Fig. 4B). These changes induced by the loss of *mep-1* in *spr-5; met-2* mutants are similar to what we observe when *lin-35* is knocked down in *spr-5; met-2* mutants. Also, similar to what we observed with knocking down *lin-35,* knocking down *mep-1* in *spr-5; met-2* mutants appears to be synergistic because knockdown of *mep-1* by itself only results in a 0.2-fold increase and a -0.1-fold decrease in the genes that are significantly misregulated in *spr-5; met-2* mutants (Fig. 4A, B).

**Figure 4.**
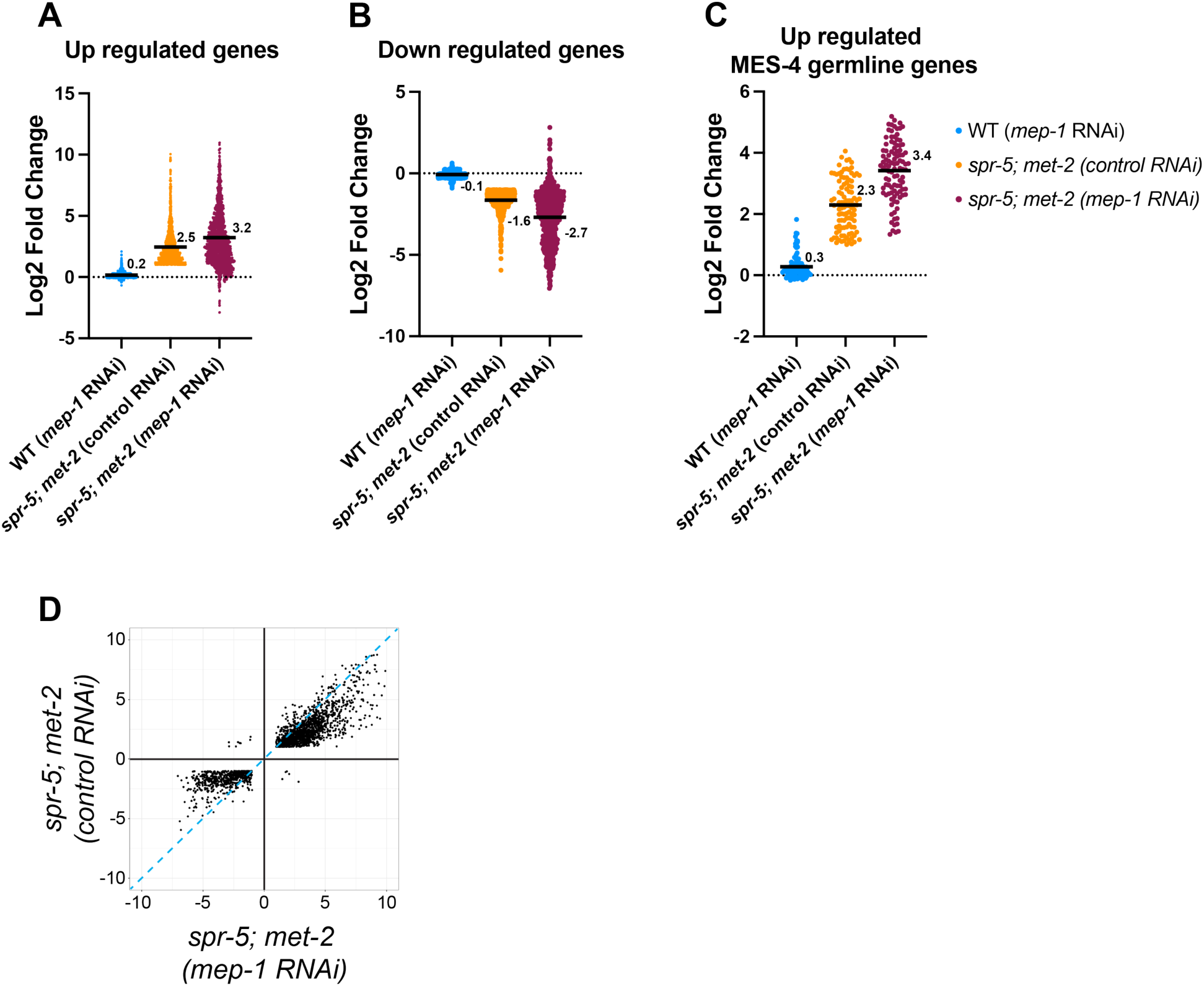
Knocking down *mep-1* further exascerbates differentially expressed genes in the soma of *spr-5; met-2* mutants. Scatter dot plots displaying the log2 fold change of the 1806 up-regulated genes (A), 895 down-regulated genes (B), and 102 up-regulated MES-4 germline genes (C) in *spr-5; met-2* mutants fed control RNAi (orange). The average log2 fold change of these same genes were examined in wild type (WT) hermaphrodites fed *mep-1* RNAi (blue) and *spr-5; met-2* mutants fed *mep-1* RNAi (maroon). (A-C) Numbers and solid black lines represent the mean log2 fold change. (D) Scatter plot displaying the correlation in log2 fold change of the all overlapping DEGs between spr-5; *met-2* mutants fed control RNAi and spr-5; *met-2* mutants *mep-1* RNAi L1 progeny. Dotted-blue line represents a one to one correlation in log2 fold change.

To determine if knocking down *mep-1* specifically exacerbates the ectopic expression of germline genes, we also compared the gene expression changes specially in the 176 MES-4 germline genes. The average log2 fold change of up regulated MES-4 germline genes in *spr-5; met-2* mutants compared to wild type is 2.3 (Fig. 4C). When *mep-1* is knocked down in *spr-5; met-2* mutants, this log2 fold change increases from 2.3 to 3.4, suggesting that the MEC NuRD complex is also reinforcing SPR-5/MET-2 repression of MES-4 germline genes in the soma (Fig. 4C). The changes in MES-4 germline gene expression appear to be synergistic because knocking down *mep-1* by itself only results in a 0.3-fold increase of the MES-4 germline genes.

To examine the overall specificity of the interaction between the MEC NuRD complex and SPR-5/MET-2 reprogramming, we also examined the gene expression changes of all 2,150 number of genes that were changed in both *spr-5; met-2* mutants and *spr-5; met-2* mutants fed *mep-1* RNAi. 99.4% were changed in the same direction and of these, 86.8% were exacerbated in *spr-5; met-2* mutants fed *mep-1* RNAi compared to *spr-5; met-2* mutants alone. Only 13 genes were in an anti-correlated quadrant (Fig. 4D). Taken together, these results suggest that, like we observe for the DREAM complex, the MEC NuRD complex is functioning specifically to reinforce SPR-5/MET-2 maternal reprogramming.

### The DREAM and MEC NuRD complexes directly bind to SPR-5/MET-2 targets

Our combined developmental delay and transcriptomics results are consistent with a model where the DREAM and MEC NuRD complexes function in the soma to reinforce SPR-5/MET-2 germline reprogramming. To determine whether the DREAM and MEC NuRD complexes bind directly to SPR-5/MET-2 germline reprogramming targets, we reanalyzed published LIN-35 and MEP-1 ENCODE chromatin immunoprecipiatation (ChIP) datasets (see Material and Methods). As with our RNAseq experiments, these ChIP experiments were performed on L1 larvae that only have 2 germ cells, enabling us to examine binding in the soma. Initially, we chose to examine the 2,698 genes that are significantly up regulated and the 1,556 genes that are significantly down regulated in *spr-5; met-2* mutants with *lin-35* RNAi compared to wild type. LIN-35 ChIP analysis of these targets demonstrated binding of LIN-35 just upstream of the transcription start site of genes significantly up regulated in *spr-5; met-2* mutants with *lin-35* RNAi compared to wild type (Fig. 5A). This enrichment is greatly reduced in genes that are significantly down regulated in *spr-5; met-2* mutants with *lin-35* RNAi compared to wild type, suggesting that the DREAM complex predominantly acts as a repressor (Fig. 5B). The binding of LIN-35 to misregulated targets appears to be specific, since we do not detect any enrichment in a set of 2,699 randomly chosen genes (Fig. 5C). Surprisingly, amongst the binding of LIN-35 to the 2,698 significantly up regulated genes, LIN-35 does not seem to bind to the genes that are most highly up regulated (boxed in Fig. 5A). It is unclear why this is the case, but it suggests that these most highly up regulated genes are indirectly affected.

**Figure 5.**
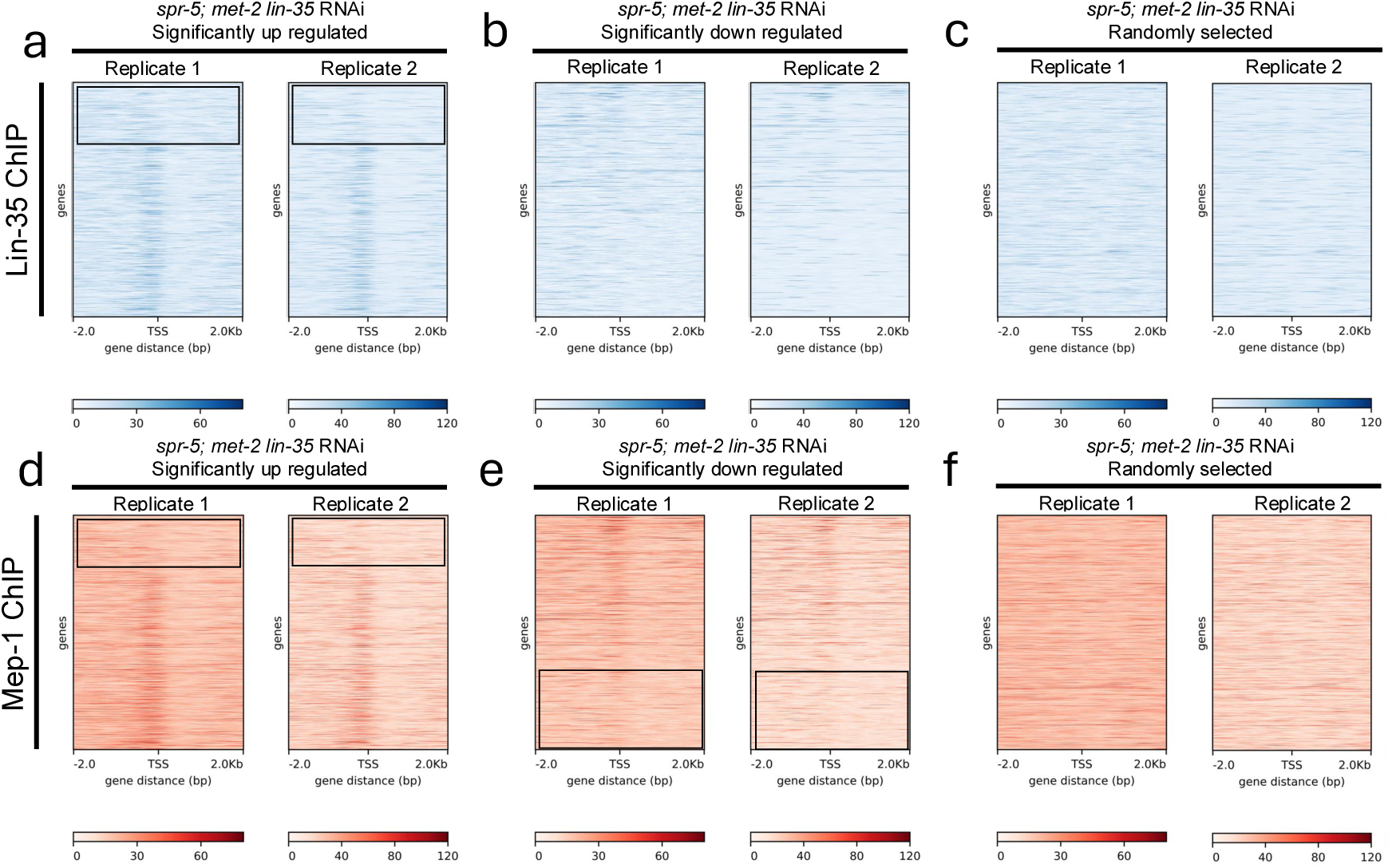
Re-analysis of LIN-35 and MEP-1 ChIP-seq data performed on LIN-35 targets exaserbated in *spr-5; met-2* mutants. (A-F) Two replicate heat maps aligned from -2.0Kb to +2.0Kb with respect to the transcriptional start site (TSS) of LIN-35 (A-C) and MEP-1 binding (D-F) show a similar enrichment (darker blue or darker red in the heat maps) of binding around the TSS at genes that are significantly up regulated (A, D) upon knockdown of *lin-35* in *spr-5; met-2* mutants (from Fig. 3A), compared to no enrichment at an identical number of randomly chosen genes (C, F). The enrichment of LIN-35 binding is completely gone at genes down regulated upon knockdown of *lin-35* in *spr-5; met-2* mutants (B), suggesting that LIN-35 acts predominantly as a repressor. In contrast, the enrichment of MEP-1 at genes down regulated upon knockdown of *lin-35* in *spr-5; met-2* mutants (from Fig. 3B) is reduced but not completely eliminated, suggesting that MEP-1 may sometimes function as an activator. In A-C the genes are listed from most highly up regulated at the top to least highly up regulated at the bottom. In D-F, the genes are listed from least highly down regulated at the top to most highly down regulated at the bottom. The boxes indicate the most highly up regulated (A, D) or down regulated genes (E) which do no bind LIN-35 (A) or MEP-1 (D, E). Similar results were obtained for analysis of genes up regulated and down regulated upon knockdown of *mep-1* in *spr-5; met-2* mutants (see supplementary Fig. 3).

To determine whether the MEC NuRD complex binds directly to SPR-5/MET-2 germline reprogramming targets in the soma, we also reanalyzed published MEP-1 ChIP data performed on L1 larvae (see Material and Methods). Examination of the 2,698 genes that are significantly up regulated in *spr-5;met-2* mutants with *lin-35* RNAi compared to wild type demonstrated an enrichment for binding of MEP-1 just up stream of the transcription start site (Fig. 5D). Since the MEC NuRD complex is thought to predominantly act as a repressor, we expected to see much less enrichment of MEP-1 binding to genes that are significantly down regulated in *spr-5; met-2* mutants with *lin-35* RNAi compared to wild type. We do observe less enrichment of MEP-1 at down regulated genes, but we still detect MEP-1 binding just upstream of the transcriptional start site in genes that are down regulated in *spr-5; met-2* mutants fed *lin-35* RNAi (Fig. 5E). This suggests that the MEC NuRD complex may sometimes be functioning as an activator in *C. elegans*. As with DREAM, the binding of MEP-1 appears to be specific, since we do not detect any enrichment in a set of 2,698 randomly chosen genes (Fig. 5F). Similar to what we observed with DREAM, MEP-1 also does not seem to bind to the genes that are most highly up regulated nor to the genes that are the most highly down regulated (boxed in Fig. 5D,E). It is unclear why this is the case, but it is consistent with these most highly changed genes being indirect.

It is possible that the DREAM and MEC NuRD complexes predominantly function together at SPR-5/MET-2 germline reprogramming targets. Alternatively, the DREAM and MEC NuRD complexes may be binding and repressing distinct subsets of SPR-5/MET-2 germline reprogramming targets. To distinguish between these possibilities, we compared the LIN-35 (Fig. 5A) and MEP-1 (Fig. 5D) ChIP data for the genes significantly up regulated in *spr-5; met-2* mutants with *lin-35* RNAi. The pattern of LIN-35 and MEP-1 binding to these genes is strikingly similar, suggesting that the DREAM and MEC NuRD complexes predominantly work together to repress SPR-5/MET-2 germline reprogramming targets.

To confirm this overlap, we also examined LIN-35 and MEP-1 binding to the 2,857 genes that are significantly up regulated and the 2,317 genes that are significantly down regulated in *spr-5;met-2* mutants with *mep-1* RNAi compared to wild type. This analysis revealed one slight difference in the binding of LIN-35 to down regulated MEC NuRD targets (Supplementary Fig. 3B) versus the binding of LIN-35 to down regulated DREAM targets (Fig. 5B). Similar to what we obaserve for MEP-1 binding (Fig. 5E and Supplementary Fig. 3E), we detect slight LIN-35 binding to the least down regulated MEC NuRD targets (Supplementary Fig. 3B). This hints that, like the MEC NuRD complex, DREAM may also ocassionally function as an activator. Nevertheless, overall the binding of both LIN-35 and MEP-1 to MEC NuRD targets is highly similar to each other (Supplementary Fig. 3) and highly similar to the binding of LIN-35 and MEP-1 to DREAM targets (Fig. 5 vs. Supplementary Fig. 3). This further suggests that the DREAM and MEC complexes predominantly work together to repress SPR-5/MET-2 germline reprogramming targets.

## DISCUSSION

Tissue specific gene expression can be influenced at a number of different levels. Of particular interest is how transcriptional repressor complexes, ATP dependent chromatin remodeling complexes and histone modifications are coordinately regulated to achieve the desired tissue specific gene expression. However, a mechanistic understanding of how these three levels of gene regulation are coordinated has been difficult to decipher. To better understand this interaction, we examined how these gene regulatory mechanisms function together to help specify the germline versus soma in *C. elegans*.

Previously we showed that when maternal epigenetic reprogramming is compromised in an *spr-5; met-2* double mutant, there is a developmental delay at the L2 larval stage in the progeny caused by the inappropriate expression of germline genes in the soma (Carpenter et al., 2021). To determine how the DREAM transcriptional repressor complex and the MEC NuRD ATP-dependent remodeling and histone deacetylation complex interact with maternal SPR-5/MET-2 epigenetic reprogramming, we asked how knocking down members of each of these complexes affects the developmental delay and ectopic germline gene expression induced by the loss of SPR-5 and MET-2. We find that loss of either DREAM or the MEC NuRD complex converts the *spr-5; met-2* L2 larval delay into an L1 larval arrest. This suggests a genetic interaction in which DREAM and the MEC NuRD complex reinforce SPR-5/MET-2 maternal epigenetic reprogramming. To determine whether the genetic interaction between these different levels of gene regulation is specific, we performed RNAseq on L1 larvae to examine the ectopic expression of germline genes in somatic tissues. This analysis demonstrated that loss of either the DREAM or MEC NuRD complexes specifically exacerbates the ectopic expression of germline genes in somatic tissues caused by loss of SPR-5 and MET-2. This suggests that the DREAM and MEC NuRD complexes specifically reinforce SPR-5/MET-2 maternal epeigenetic reprogramming in the soma.

The DREAM and MEC NuRD compexes may function directly at SPR-5/MET-2 maternal reprogramming targets or indirectly through other genes. To distinguish between these possibilities, we re-analyzed previously published DREAM and MEC NuRD complex ChIP data. Both the DREAM and MEC NuRD complexes are enriched at SPR-5/MET-2 maternal reprogramming targets, suggesting they act directly on SPR-5/MET-2 maternal reprogramming targets. It is possible that the Dream and MEC NuRD complexes are individually acting on mutually exclusive subsets of SPR-5/MET-2 maternal reprogramming targets. Alternatively, the DREAM and MEC NuRD compexes may both be functioning together at SPR-5/MET-2 maternal reprogramming targets. To distinguish between these possibilities, we compared the ChIP binding profiles of the DREAM and MEC NuRD compex members to one another at SPR-5/MET-2 maternal reprogramming targets. The binding profiles of the DREAM and MEC NuRD compex members to SPR-5/MET-2 maternal reprogramming targets are remarkably similar, suggesting that they act together directly at SPR-5/MET-2 maternal reprogramming targets. The high degree of overlap in binding of LIN-35 and MEP-1 at SPR-5/MET-2 maternal reprogramming targets also raises the possibility that one of these complexes may be recruiting the other. For example, the DREAM complex may predominantly function to recruit the MEC NuRD complex. However, the function of the DREAM and MEC NuRD complexes at these targets does not appear to be redundant because loss of either the DREAM or MEC NuRD complex members is sufficient on their own to exacerbate the developmental delay phenotype.

### A model for how the maternal reprogramming of histone methylation is reinforced by transcriptional repression and ATP-dependent chromatin remodeling

We propose that the DREAM and MEC NuRD complexes reinforce SPR-5/MET-2 maternal reprogramming through the following model. H3K4me2 is acquired during transcription in the germline and functions as a transcriptional memory helping to maintain transcription of germline genes (Lee and Katz, 2020). SPR-5 and MET-2 function at fertilization as a maternal epigenetic reprogramming switch to remove the H3K4me2 transcriptional memory and replace it with with H3k9me2 to further repress these genes (Andersen and Horvitz, 2007; Greer et al., 2014; Katz et al., 2009; Kerr et al., 2014). SPR-5/MET-2 maternal reprogramming is necessary to reset the entire genome to a ground state and prevent the H3K4me2 transcriptional memory from being inappropriately propagated to the next generation where it could result in the ectopic expression of germline genes in the embryo.

In the embryo of the next generation, the germline blastomere P4 is specified after just four cell divisions. H3K36me3, which is acquired during transcription in the germline via the transcription dependent H3K36 methyltransferase MET-1, is maintained in the embryo by a transcription independent H3K36 methyltransferase MES-4 (Furuhashi et al., 2010; Kreher et al., 2018; Rechtsteiner et al., 2010). This maintained H3K36me3 “bookmark” prevents the chromatin at germline genes from being repressed by maternal SPR-5/MET-2 reprogramming and facilitates the re-initiation of germline transcription to help specify the germline. The SPR-5/MET-2 maternal reprogramming and MES-4 bookmarking systems are mutually antagonistic, because in the absence of SPR-5/MET-2 maternal reprogramming, MES-4 inappropriately maintains H3K36me3 at germline genes in the soma. This results in the expression of germline genes in the soma (Carpenter et al., 2021).

Our work here suggests that the initiation phase of germline versus soma distinction, facilitated by SPR-5/MET-2 and MES-4, is then reinforced in the soma by the DREAM and MEC NuRD complexes. As the soma differentiates during late embryonic and larval development, transcription factors (TFs) drive tissue specific gene expression. To prevent cross talk between these tissue specific TFs and germline genes which could result in the inappropriate expression of germline genes in somatic tissues, the repression of chromatin at germline genes that was initiated by SPR-5/MET-2 reprogramming in the initiation phase is reinforced in the soma during a maintenance phase by the DREAM and MEC NuRD complexes. This function of the DREAM and MEC NuRD complexes during the maintenance phase is consistent with previous data showing that MEP-1 and LIN-35 have defects in germline versus soma specification (Rechtsteiner et al., 2019; Unhavaithaya et al., 2002; Wang et al., 2005; Wu et al., 2012).

### Why is transcriptional repression and ATP-dependent chromatin remodeling required in the soma to reinforce SPR-5/MET-2 maternal epigenetic reprogramming?

In the very early *C. elegans* embryo, there is no transcription (Baugh et al., 2003; Edgar et al., 1994; Seydoux and Dunn, 1997). Even after transcription begins in the *C. elegans* embryo at the 8-16 cell stage, the bulk of transcriptional elongation does not occur until the ∼60 cell stage. As a result, the *C. elegans* embryos can survive until the 100 cell stage without RNA polymerase II (RNA pol II) because embryogenesis runs on maternally contributed factors (Powell-Coffman et al., 1996). Because of this lack of transcription in the early *C. elegans* embryo, the loss of H3K4me2 and addition of H3K9me2 may be sufficient in the early embryo to establish a repressed state genome wide. However, during later embryogenesis and the larval stages, somatic tissues differentiate and have significant tissue specific gene expression driven by tissue specific transcription factors (Murray et al., 2008). Because the tissue specific TF binding sites may overlap slightly with the promoters of germline genes, there may be a more stringent requirement to block gene expression at multiple levels at these later stages; including via ATP dependent chromatin remodeling, histone deacetylation and trancriptional repression. This could potentially be why the DREAM and MEC NuRD are required in the soma to reinforce SPR-5/MET-2 maternal epigenetic reprogramming during the maintenance phase of germline versus soma specification.

Our data demonstrate that SR-5/MET-2 maternal reprogramming is not sufficient to fully repress germline transcription in the soma. But how do the DREAM and MEC NuRD complexes provide additional gene repression that is distinct from loss of H3K4me2 and acquisition of H3K9me2? DREAM is thought to function as a transcriptional repressor, but does not have any known enzymatic activity associated with the complex (Hoareau et al., 2024). Thus it may function by binding to promoters and specifically competing against TF binding and RNA Pol II binding to prevent transcriptional initiation. Alternatively, or in addition, the DREAM complex may recruit the MEC NuRD complex, but this remains to be tested. The MEC NuRD complex has both ATP-dependent chromatin remodeling activity and HDAC activity associated with it (Xue et al., 1998). The ATP-dependent chromatin remodeling activity of NuRD and the HDAC activity associated with the NuRD complex are thought to function by restricting nucleosome spacing and tightening the association between nucleosomes and DNA (Denslow and Wade, 2007). It is not entirely clear why ATP dependent chromatin remodeling, histone deacetylation and transcriptional repression are needed on top of the addition of the loss of H3K4me2 and addition of H3K9me2. However, our finding that loss of either DREAM or the MEC NuRD complex exacerbates the *spr-5; met-2* double mutant clearly indicates that the loss of H3K4me2 and addition of H3K9me2 are not functionally equivalent to gene repression via ATP dependent chromatin remodeling, histone deacetylation and trancriptional repression. Thus, our work gives insight into the relationship between histone modifications, nucleosome remodeling and transcriptional repressors. Specifically, it suggests that to fully repress transcription, it may be necessary to block transcription at each of these levels.

## Acknowledgements

NuRD and DREAM complex RNAi enhancer screens were piloted by undergraduate students at Oglethorpe University as part of the *C. elegans* Pipeline CURE embedded in their biology major. The research cohort included 114 students (68 students from Developmental Biology (BIO 313 - SP 2018, SP 2020 and SP2022) and 46 students from Research in Epigenetics (BIO 455 - FA 2018, FA 20219, FA 2021 and FA 2023)). The Pipeline CURE supported and continues to support the Broader Impact goals of NSF Grants IOS1354998, IOS 1931697 and IOS2418128. We thank the *Caenorhabditis* Genetics Center (funded by NIH P40 OD010440) for strains and WormBase (WS292).

## Data Availability Statement

Raw and processed RNAseq files have been deposited into Gene Expression Omnibus (www.ncbi.nlm.nih.gov/geo) under accession code GSE303062 (*lin-35* RNAi RNAseq) and GSE303085 (*mep-1* RNAi RNAseq).

---

**Supplemental Figure 1.**
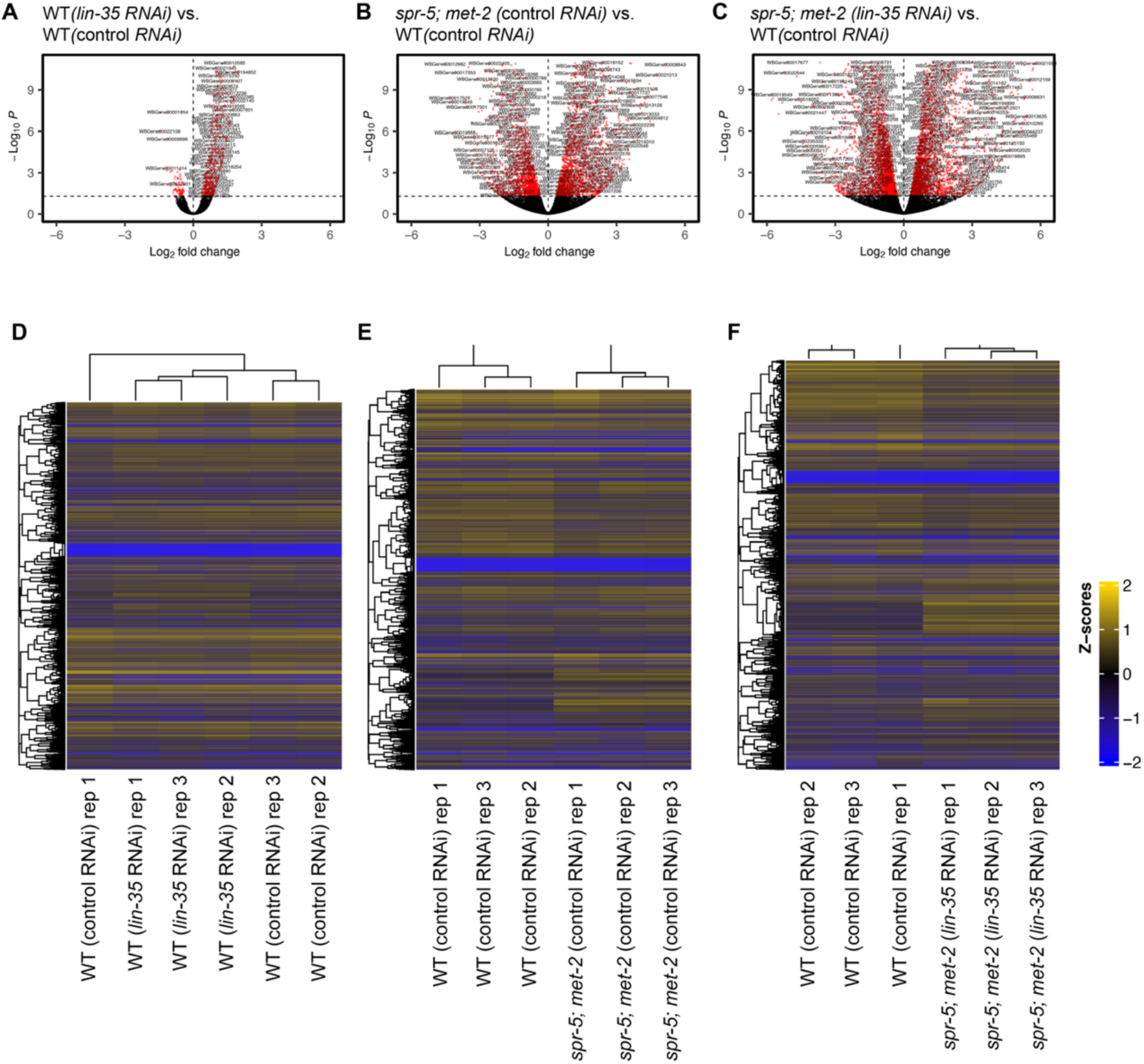
: Differential expression and replicate comparison of RNAseq experiments performed on wild type, *spr-5; met-2* progeny fed either control or *lin-35* RNAi. Volcano plot of log2 fold changes in gene expression (x-axis) by statistical significance (-Log_10_ P-value; y-axis) in L1 progeny of wild type (WT) hermaphrodites fed *lin-35* RNAi (A), *spr-5; met-2* hermaphrodites fed L4440 (control) RNAi (B), and *spr-5; met-2* hermaphrodites fed *lin-35* RNAi (C) compared to L1 progeny of wild type (WT) hermaphrodites fed control RNAi. Heatmap of differentially expressed RNA-seq transcripts between L1 progeny of wild type (WT) hermaphrodites fed control RNAi and wild type (WT) hermaphrodites fed *lin-35* RNAi (D), *spr-5; met-2* fed L4440 (control) RNAi (E), and *spr-5; met-2* fed *lin-35* RNAi (F). Data was scaled and hierarchical clustering was performed using complete linkage algorithm, with distance measured by calculating pairwise distance. Higher (blue) and lower (yellow) expression is reported as a z-score.

**Supplemental Figure 2.**
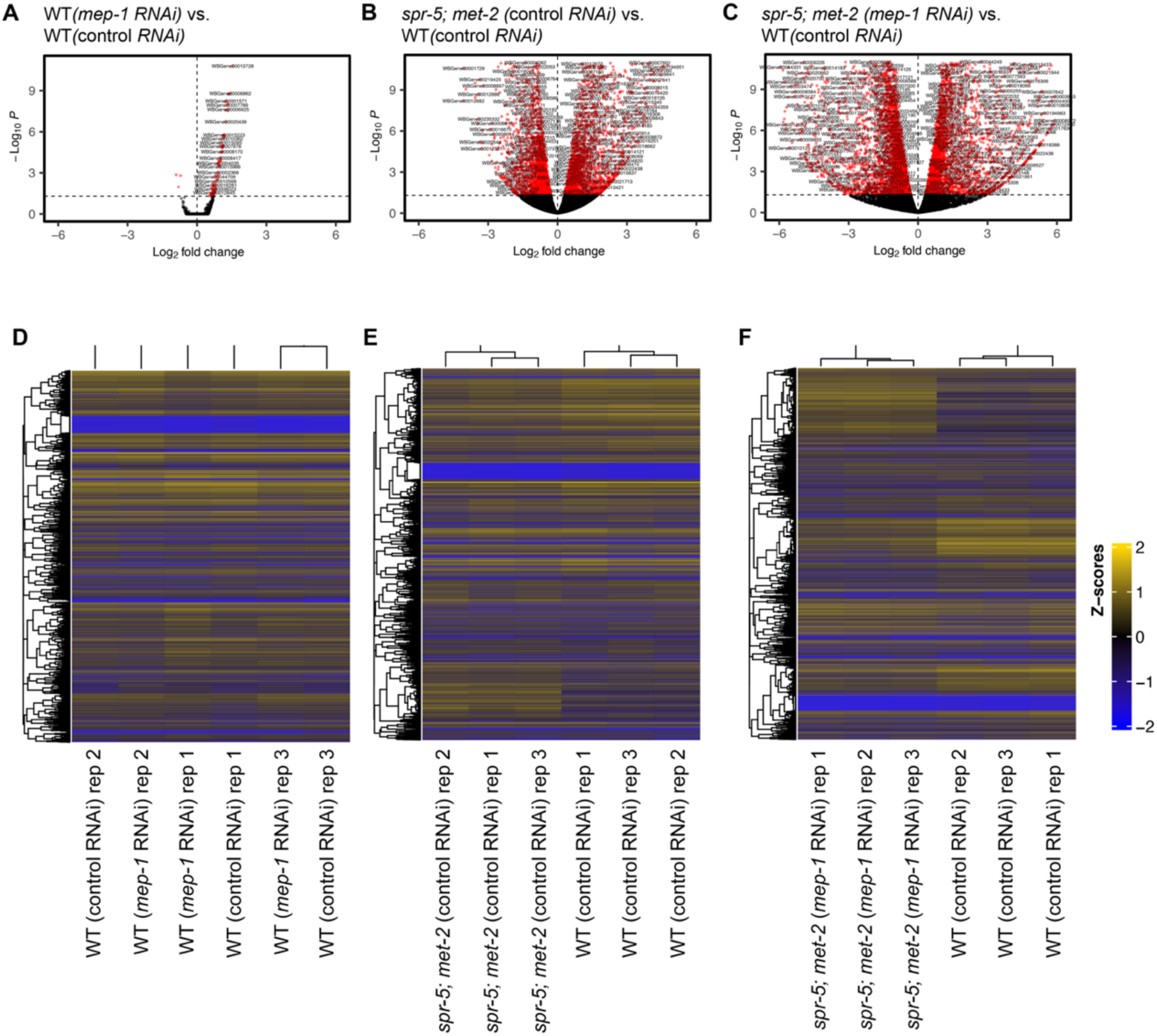
: Differential expression and replicate comparison of RNAseq experiments performed on wild type, *spr-5; met-2* progeny fed either control or *mep-1* RNAi. Volcano plot of log2 fold changes in gene expression (x-axis) by statistical significance (-Log_10_ P-value; y-axis) in L1 progeny of wild type (WT) hermaphrodites fed *mep-1* RNAi (A), *spr-5; met-2* hermaphrodites fed L4440 (control) RNAi (B), and *spr-5; met-2* hermaphrodites fed L4440 *mep-1* RNAi (C) compared to L1 progeny wild type (WT) hermaphrodites fed control RNAi. Heatmap of differentially expressed RNA-seq transcripts between L1 progeny of wild type (WT) hermaphrodites fed control RNAi and wild type (WT) hermaphrodites fed *mep-1* RNAi (D), *spr-5; met-2* fed L4440 (control) RNAi (E), and *spr-5; met-2* fed *mep-1* RNAi (F). Data was scaled and hierarchical clustering was performed using complete linkage algorithm, with distance measured by calculating pairwise distance. Higher (blue) and lower (yellow) expression is reported as a z-score.

**Supplemental Figure 3.**
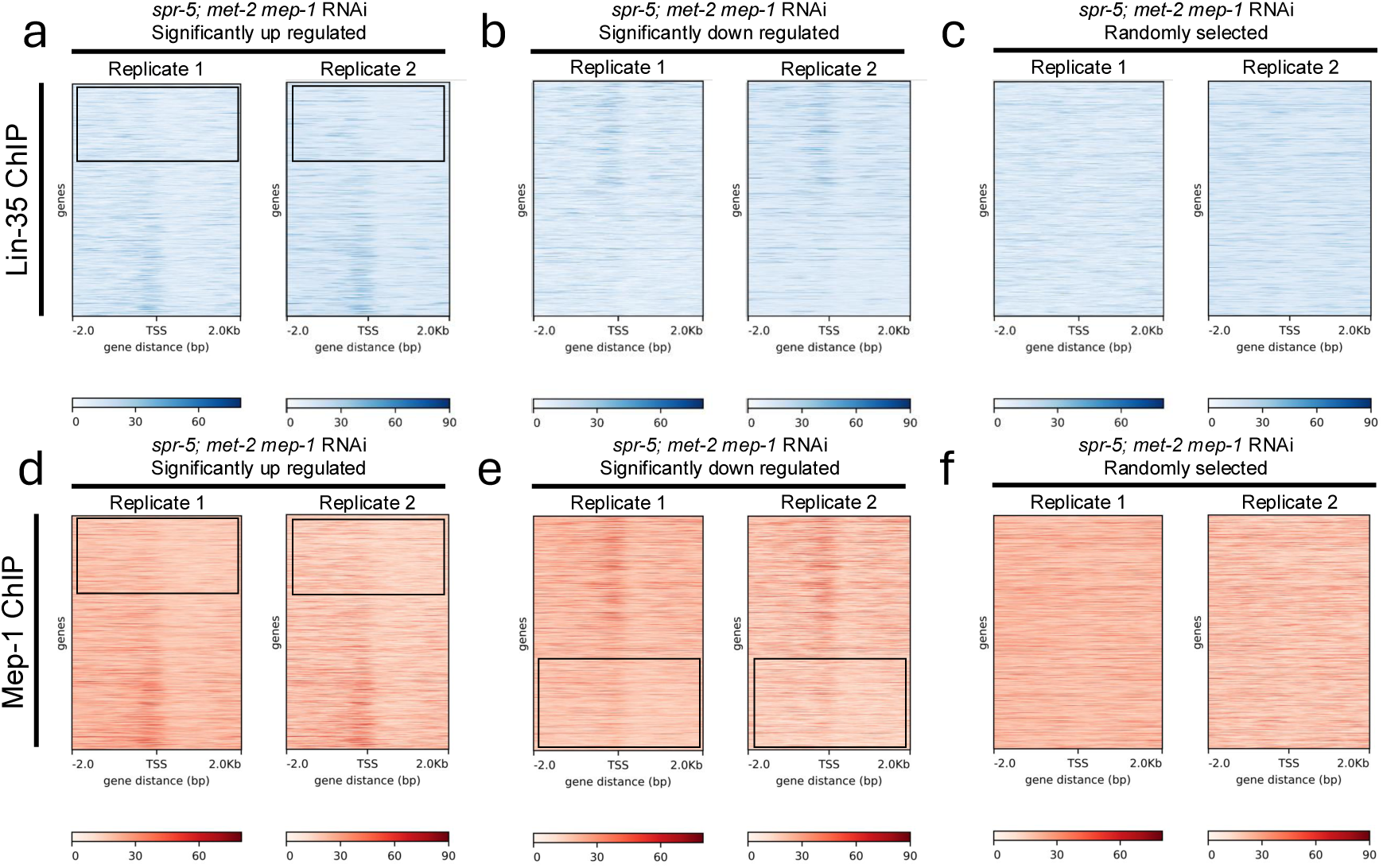
: Re-analysis of LIN-35 and MEP-1 ChIP-seq data performed on MEP-1 targets exaserbated in *spr-5; met-2* mutants. (A-F) Two replicate heat maps aligned from -2.0Kb to +2.0Kb with respect to the transcriptional start site (TSS) of LIN-35 (A-C) and MEP-1 binding (D-F) show a similar enrichment (darker blue or darker red in the heat maps) of binding around the TSS at genes that are significantly up regulated (A, D) upon knockdown of *mep-1* in *spr-5, met-2* mutants (from Fig. 4A), compared to no enrichment at an identical number of randomly chosen genes (C, F). The enrichment of LIN-35 binding is far less, but not completely gone at genes down regulated upon knockdown of *lin-35* in *spr-5, met-2* mutants (B) (from Fig. 4B), suggesting that LIN-35 acts predominantly as a repressor, but may ocassionally function as an activator. The enrichment of MEP-1 at genes down regulated upon knockdown of *lin-35* in *spr-5, met-2* mutants (from Fig. 4B) is reduced but not completely eliminated, suggesting that MEP-1 may sometimes function as an activator. In A-C the genes are listed from most highly up regulated at the top to least highly up regulated at the bottom. In D-F, the genes are listed from least highly down regulated at the top to most highly down regulated at the bottom. The boxes indicate the most highly up regulated (A, D) or down regulated genes (E) which do no bind LIN-35 (A) or MEP-1 (D, E).

